# European agroforestry is no universal remedy for biodiversity: a time-cumulative meta-analysis

**DOI:** 10.1101/2020.08.27.269589

**Authors:** Anne-Christine Mupepele, Matteo Keller, Carsten F Dormann

## Abstract

**Background:** Agroforestry is a production system combining trees with crops or livestock. It has the potential to increase biodiversity in relation to single-use systems, such as pastures or conventional agriculture, by providing a higher habitat heterogeneity. In a literature review and subsequent meta-analysis, we investigated the relationship between biodiversity and agroforestry and critically appraise the underlying evidence of the results.

**Results:** Biodiversity in agroforestry was higher than in conventional agriculture, but could not outcompete pastures, forests and abandoned agroforestry systems. There was no overall biodiversity benefit in agroforestry systems. Data were available for plants, birds, bats and arthropods. Arthropods and birds were the two taxonomic groups profiting from agroforestry systems. A time-cumulative meta-analysis shows that there was no general benefit of biodiversity at any point in the past besides in early 2015. Time-cumulative meta-analysis can unravel missing robustness of meta-analytical results if conclusions alternate between significant to non-significant summary effect sizes over time.

**Conclusion:** Agroforestry increases biodiversity only in silvoarable systems compared with conventional agriculture. But even this result is based on a small magnitude and single-study effect sizes were heterogeneous with sometimes opposing conclusions. The latter suggests the importance of other usually unmeasured variables, such as landscape parameters or land-use history, influencing biodiversity in agroforestry systems.

## Introduction

Agroforestry is a collective name for diverse land-use systems integrating tree husbandry with livestock or arable cultivation [1, 2]. It is a key historical element of the European landscape currently experiencing changes from traditional systems, e.g. large fruit tree orchards with extensive livestock grazing, to newer approaches, e.g. short rotation coppice in combination with crop rows [3]. Agroforestry is classified into silvopastoral systems, grazed by livestock or used for fodder production, and silvoarable systems, in which crops are grown among trees [4]. Fields where trees are grown only at the edge, such as stream side management zones or hedgerows adjacent to arable land, are also occasionally subsumed under agroforestry systems [4]. In these cases, the herbaceous and wooded components are usually not managed together and often have different owners. In our study, trees or shrubs adjacent to fields or pastures are not considered.

Biodiversity is threatened and particularly steep declines have been observed in intensively used agricultural areas [5–8]. Compared to monocultures, agroforestry systems increase heterogeneity in the landscape structure and potentially lead to increased biodiversity [5, 9–12]. Demonstrating a clear benefit for biodiversity could favour future subsidies for agroforestry systems by the Common Agricultural Policy or its successor policies [13–17]. The benefits for biodiversity in agroforestry systems have been investigated particularly in the tropics showing that biodiversity can be improved by agroforestry in degraded and intensively cropped areas [18–20]. In the temperate zones, studies for different species groups, such as birds [21] or invertebrates [22] have shown equivocal effects on biodiversity. An earlier meta-analysis found a net increase of biodiversity across taxa and agroforestry systems in Europe, however failed to provide detailed information on the heterogeneity and robustness of their findings [23]. Here we provide an evidence update and a more explicit discussion on biodiversity in comparison to forests and agriculture/pasture and assess the robustness of the results by answering the following research questions: (1) Does agroforestry affect biodiversity? (2) Is biodiversity in agroforestry influenced by other environmental variables, such as differences in taxonomic groups or climatic region? (3) How robust and strong is the underlying evidence of these results?

## Materials and Methods

We review the literature on biodiversity in European agroforestry systems and synthesize the results in a meta-analysis. This review is based on the standards of the Collaboration for Environmental Evidence [24–27]. It further goes beyond these standards by additionally performing a sensitivity analysis with studies weighted based on their evidence to identify the robustness of the results [28].

### Literature search

We used search terms and their synonyms related to ‘biodiversity’, ‘agroforestry’ and ‘Europe’ in the Web of Science to identify the relevant literature (Box 1). Reviews revealed by the Web of Science search were scanned for additional references. In the first screening of articles, we sighted title and abstract and excluded publications that did not fulfil the inclusion criteria (Box 2). In a second screening, we read the full text and applied additional inclusion criteria (Box 2). If an article was included, we extracted the mean diversity, standard deviation and sample size in an agroforestry system and its corresponding control site along with environmental variables (Table 1). WebPlotDigitizer was used to extract data points from figures [29]. Unique combinations of agroforestry system, control type and taxonomic group were considered from each article.

**Table 1:** Environmental variables.

| Variable name | Categories |
| --- | --- |
| agroforestry system | silvopastoral; silvoarable |
| control type | forest, (conventional) agriculture (i.e. pure crop fields), pasture or abandoned agroforestry systems (generally described as shrub-encroached) |
| sampling methods | transects with sweep netting; pitfall traps; pan traps; recording and various other methods |
| diversity measure | species richness, family richness or Shannon diversity |
| sampling year | numeric ranging from 1984 to 2013 |
| country of study location | European country |
| climate zone | Mediterranean (including two summer-moist Atlantic studies), temperate Central European or boreal |

#### Box 1: Search string used in the Web of Science initially in January 2016 with updates in 2018, 2019 and last on 14th February 2020. The search covered the following databases: Web of Science Core Collection, BIOSIS Citation Index, BIOSIS Previews, Current Contents Connect, Data Citation Index, Derwent Innovations Index, KCI-Korean Journal Database, MEDLINE, SciELO Citation Index, Zoological Record. Search options in the Web of Sciences were set to ‘all years’ and ‘all languages’.

Topic: (*diversity OR “species richness” OR “species composition”) AND: (Agroforest* OR agro-forest* OR silvopast* OR *silvoarabl* OR dehesa OR “alley* cropping” OR “wood* pasture*” OR “forest* farming*” OR “orchard* intercropping” OR “scatter* tree*” OR “grazed orchard” OR montado) AND: (Europe OR Albania OR Andorra OR Armenia OR Austria OR Azerbaijan OR Belarus OR Belgium OR “Bosnia and Herzegovina” OR Bulgaria OR Croatia OR Cyprus OR “Czech Republic” OR Denmark OR Estonia OR Finland OR France OR Georgia OR Germany OR Greece OR Hungary OR Iceland OR Ireland OR Italy OR Latvia OR Liechtenstein OR Lithuania OR Luxembourg OR Macedonia OR Malta OR Moldova OR Monaco OR Montenegro OR Netherlands OR Norway OR Poland OR Portugal OR Romania OR Russia OR Serbia OR Slovakia OR Slovenia OR Spain OR Sweden OR Switzerland OR Ukraine OR “United Kingdom” OR England OR Wales OR Scotland)

### Analysis

Meta-analysis is based on effect sizes and here we used log response ratios to compare the biodiversity between an agroforestry site and its corresponding control site [30, 31]. The summary effect of agroforestry on biodiversity was estimated by running a random-effect model, with a random effect on the study, and no further fixed effects [32]. Heterogeneity was tested with a Q test for heterogeneity and additionally given by *I*^2^, the ratio of heterogeneity (i.e. between-study variability) to the total variability (i.e. sum of between- and within-study variability) [33, 34]. If heterogeneity accounts for large amounts of the total variability, additional environmental variables (moderators) may improve the model by further explaining parts of the heterogeneity. This was investigated with a mixed-effects model with fixed-effects selection based on a likelihood-ratio test [35]. Marginal *R*^2^ was given to identify the amount of heterogeneity that could be explained by the selected mixed-effects model [36, 37]. If the mixed-effects model identified categorical environmental variables influencing the agroforestry-biodiversity relationship, a subsequent subgroup meta-analysis was performed to identify under which circumstances agroforestry is beneficial to biodiversity. Analysis was realized in R 4.0.2 using packages ‘metafor’, ‘nlme’ and ‘MuMIn’ [30, 38, 39, see Additional file 5 for details and R code].

#### Box 2: Inclusion criteria for studies to be included in this review.

**Inclusion criteria for title and abstract screening**

- Study sites are located in Europe. Europe ranges from Iceland to the Ural Mountains and from Norway to the Mediterranean Sea and the Black Sea.

- The study is done in an agroforestry system, whereas agroforestry is defined by an area covered by crops or livestock and trees in an *alternating* way. Buffer strips and hedgerows only bordering an agricultural field or pasture were not considered.

- The study provides information on biodiversity in an agroforestry system.

**Additional inclusion criteria for full-text screening**

- The article should be accessible through the subscriptions hold by the University of Freiburg or personal communication with the authors.

- The study should not discuss conceptual approaches or introduce new methods without quantifying biodiversity in agroforestry.

- If a study reviewed other primary studies, references were verified for inclusion.

- Average species richness or another quantifiable biodiversity measure, such as Shannon diversity, needs to be extractable for an agroforestry system and a corresponding control type in relation to their sample size.

### Sensitivity analysis

Studies are traditionally weighted according to their inverse variance. This method has been criticized for being prone to bias especially with small sample sizes [40]. We tested the robustness of the results by adjusting the weighting by the underlying evidence of each study [compare with 41]. For this purpose studies were scaled according to their level of evidence [28]. Publication bias was assessed based on a funnel plot and an Egger’s regression test [31, 35, 42].

## Results

The literature search resulted in 1411 records from which 50 articles met all inclusion criteria (Fig. 1, Additional file 1). Unique combinations of agroforestry systems (silvoarable or silvopastoral), control types (forest, agriculture, pasture or abandoned agroforestry systems) and taxonomic groups per study led to 69 effect sizes used in the meta-analysis.

**Figure 1:**
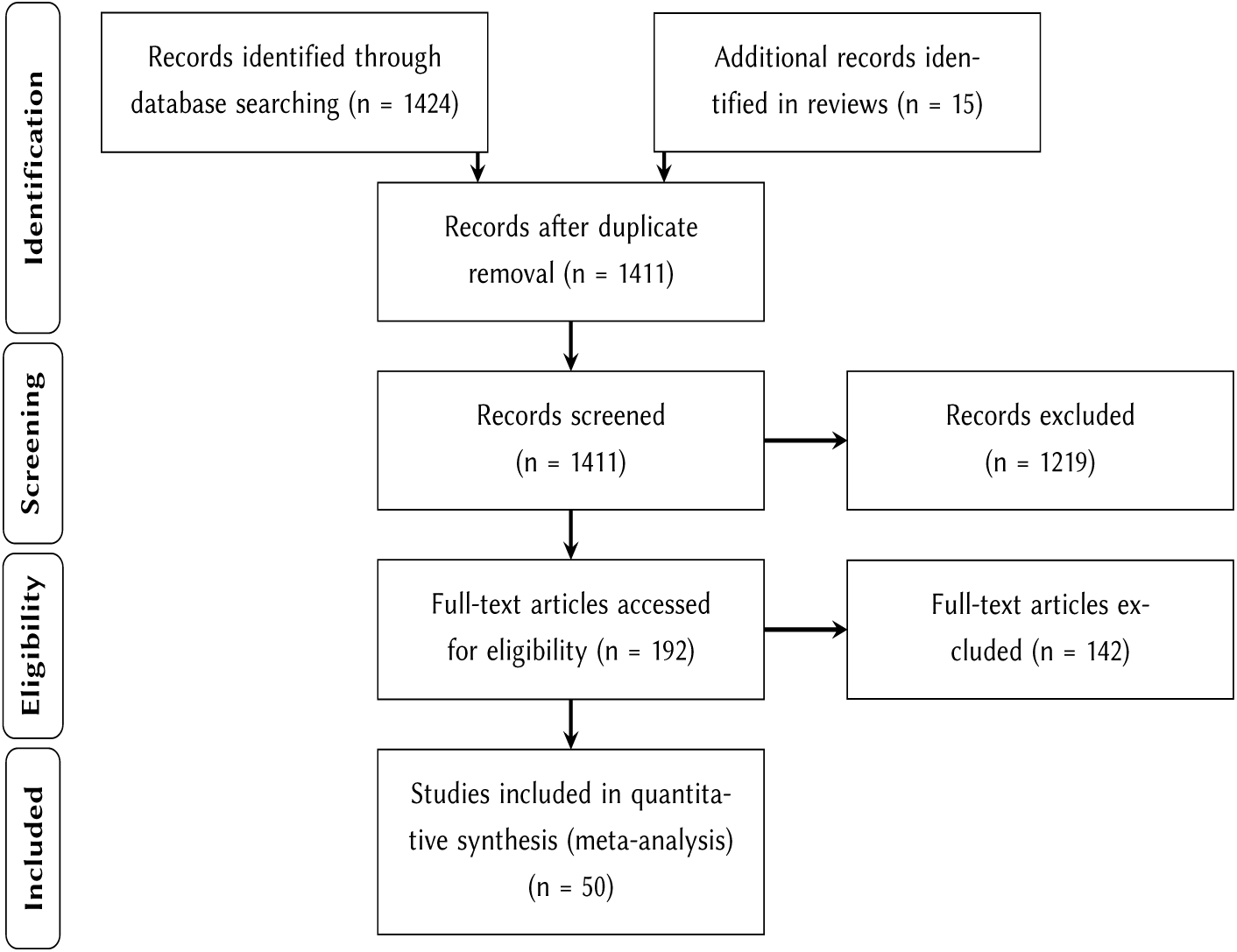
PRISMA flow diagram [25, 74].

Studies had been conducted in sites all across Europe and covered data from 1984 to 2019 (Fig. 2). The majority of study sites were located in the Mediterranean with 12 studies from Spain, 8 from Portugal, 5 from Italy and one from France and Turkey each. There were fewer studies from the temperate central European climate, characterized by cold winters and summer-green deciduous forests. They ranged from the United Kingdom (6), Romania (4), France (2), Germany (2), Switzerland (2) and Belgium (1) to northern Italy (1). The boreal region was represented by four studies from Sweden and two from Finland.

**Figure 2:**
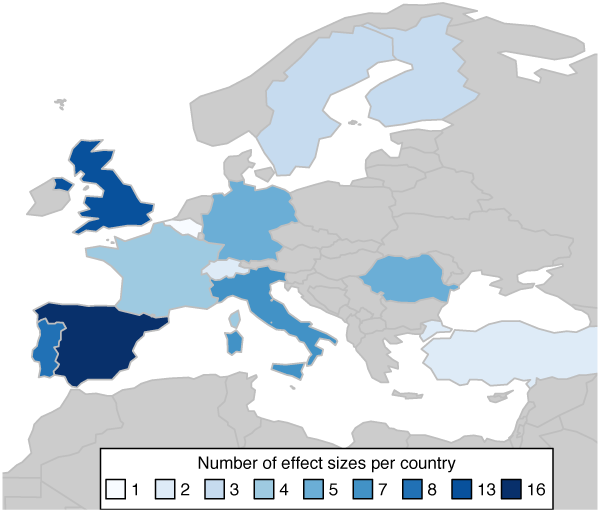
Map of Europe with the number of effect sizes per country.

**Figure 3:**
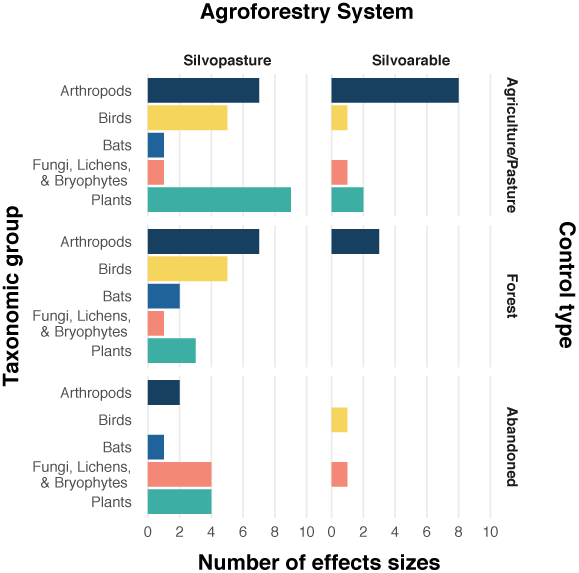
Number of effect sizes in each combination of agroforestry system, comparator and biodiversity group.

Agroforestry systems were predominately silvopastoral (36 studies, 52 effect sizes), while silvoarable systems were less often a topic of research (13 studies, 17 effect sizes). The impact of agroforestry on biodiversity was evaluated by comparing agroforestry systems to a control type. Most often this control type was a pasture (23 effect sizes), followed by forests (21 effect sizes), conventional agricultures, i.e. crop fields, (12 effect sizes) and abandoned agroforestry systems (13 effect sizes).

Biodiversity was measured in different taxonomic groups and reported at various levels of detail across studies. Some studies for example lumped all arthropods, whereas others reported diversity of carabids only. We clustered biodiversity measures into five groups: arthropods, birds, bats, plants and one group with fungi, lichen and bryophytes. Biodiversity effects were mainly measured based on differences in species richness. Five studies with seven effect sizes used other measures, namely family richness [43, 44], log-series [45, 46] or Shannon index [47].

### Effects of agroforestry on biodiversity

The results of the meta-analysis show that there is no general benefit of agroforestry systems on biodiversity (summary effect size = 0.1, 95%CI = [−0.03, 0.23], *z*_*df*=68_ = 1.47, *p* = 0.14, Additional file 5). The studies’ individual effects sizes show substantial between-study variability (Q = 6229, *p* < 0.0001; *I*^2^ = 98.9%; Fig. 4.) Some of this heterogeneity was attributed to systematic differences in environmental variables and the ‘taxonomic group’, ‘control type’ and ‘agroforestry type’ could explain 13.5% of the heterogeneity (marginal *R*^2^).

**Figure 4:**
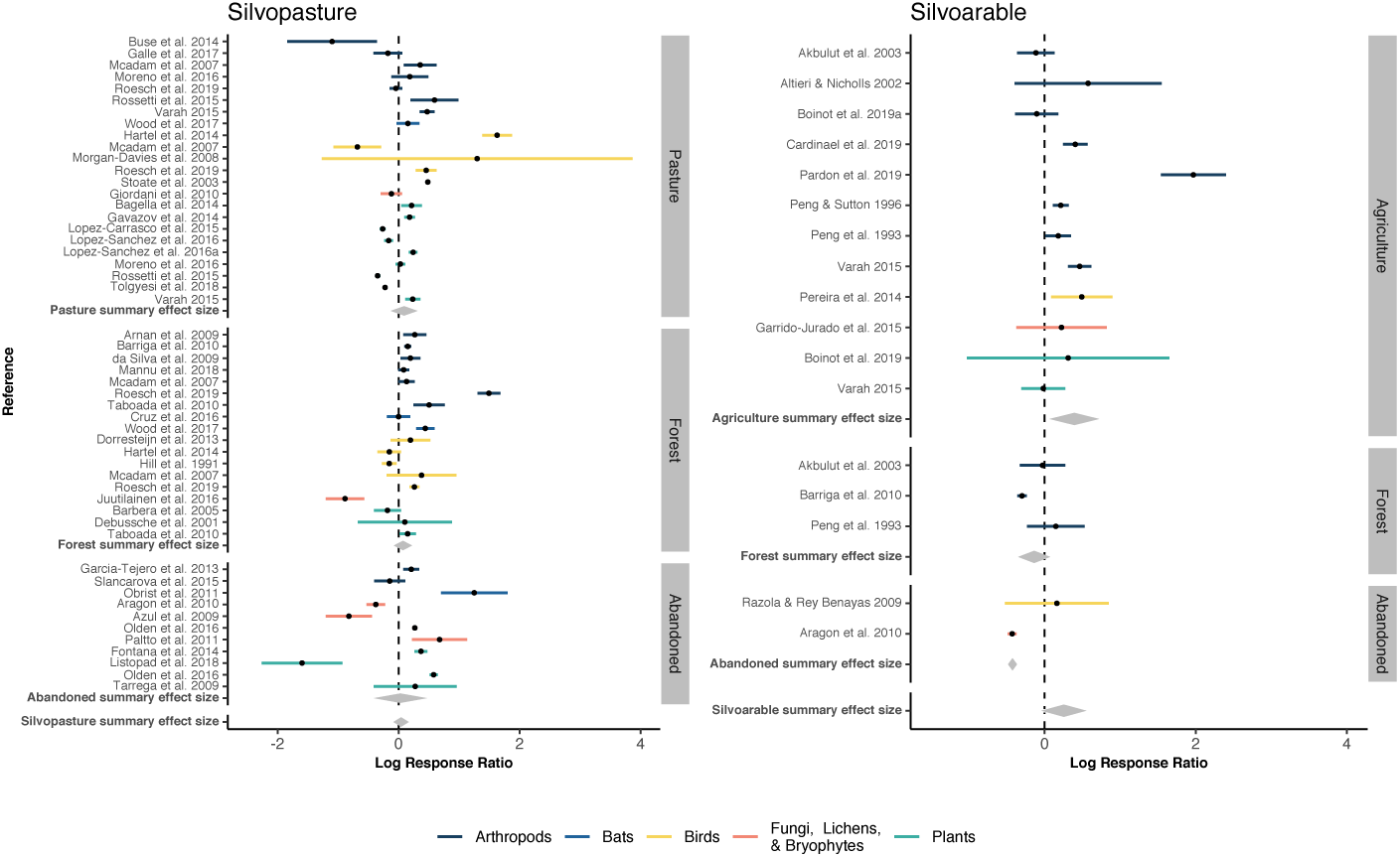
Forest plot for silvopasture and silvoarable systems with subgroup summary effect sizes (grey diamonds) per system (silvoarable, silvopastoral) and per control type (pasture/conventional agriculture, forest or abandoned agroforestry system).

A subgroup analysis for each agroforestry system, further distinguishing biodiversity effects depending on the control type, revealed that silvoarable systems were significantly more divers than conventional agriculture (Fig. 4b, ‘Agriculture’ summary effect size = 0.46, 95%CI = [0.1, 0.82], *z*_*df*=11_ = 2.52, *p* = 0.012) with 1.6 times more species in the agroforestry system than in conventional agriculture. Comparing the biodiversity of silvoarable systems to forests, they did not differ significantly, but showed a tendency towards higher diversity in forests. In silvopastoral systems none of the subgroup effect sizes was significant (Fig. 4a). Effect sizes were very heterogeneous and with partly opposing effects, such as forests harbouring a higher bird diversity in relation to agroforestry in one study [48, moderate-evidence study] and vice versa in another study [49, moderate-evidence study].

A subgroup analysis of taxonomic groups showed that birds and arthropods are significantly more diverse across all agroforestry systems (Bird summary effect size = 0.23, 95%CI = [0.012, 0.44], *z*_*df*=11_ = 2.07, *p* = 0.038; Arthropods summary effect sizem = 0.3, 95%CI = [0.016, 0.59], *z*_*df*=26_ = 2.07, *p* = 0.038). For arthropods a higher resolution was available with subgroups on different taxonomic levels, such as bees or spiders. This increased the number of effect sizes from 27 to 41 as the number of unique combination of taxonomic group and agroforestry system increased. None of the most replicated groups, i.e. beetles, bees and spiders, showed a consistent diversity response to agroforestry (Fig. 5).

**Figure 5:**
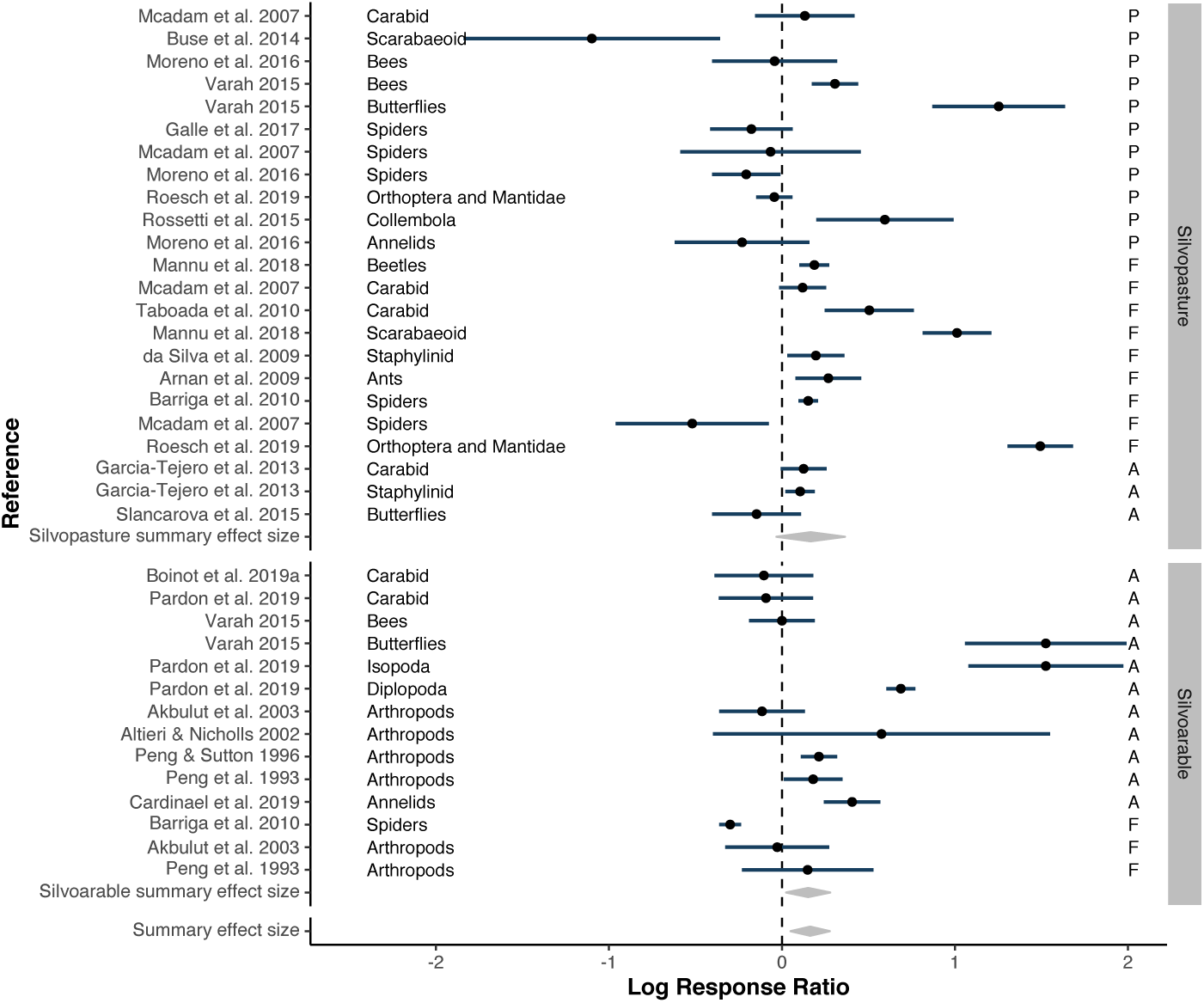
Arthropod subgroup analysis with summary effect sizes (grey diamonds). Taxonomic groups are provided in more detail depending on reported groups in primary studies. Letters on the right side reflect the first letter of the control type (A=Agriculture or in silvopastoral systems A=Abandoned, P= Pasture, F=Forests).

### Sensitivity analysis and the underlying evidence

The quality of studies included in this meta-analysis ranged from weak to strong evidence [compare with 28]. Some studies were based on a replicated and controlled design providing the strongest evidence, whereas others used before-after comparison or an observational gradient. We adjusted the study weights according to their level of evidence to assign weaker studies with a lower weight. The results of the evidence-weighted meta-analysis did not lead to different conclusions and confirmed the results of the traditional inverse-variance-weighted meta-analysis (Level-of-evidence-weighted summary effect size = 0.095, 95%CI = [−0.0063, 0.2]). Beside the weighting of studies, missing studies due to a publication bias is another obstacle for robust meta-analytical results. According to the funnel plot and Egger’s regression test, no publication bias is detectable in our data (Additional files 5 and 6, intercept of Egger’s regression = 0.77, *t* = 0.03, *p* = 0.98).

Given that an earlier meta-analysis has found a significant effect of agroforestry on biodiversity, we were interested in the change of the conclusion over time [23]. A cumulative meta-analysis shows that there is a tendency of agroforestry to be beneficial across time. But only in 2015, when the studies from Garrido-Jurado et al. [50] and Rossetti et al. [51] were added, the confidence interval was above zero (Fig. 6). A meta-analysis conducted in early 2015 would have resulted in an overall significant positive effect of agroforestry on biodiversity. During all other moments between 1991 and today, there is no general beneficial effect of agroforestry for biodiversity, and the conclusion remains robust over the time. Another possible bias could have been introduced by systematically investigating a particular taxonomic group during a certain time period, e.g. a peak of bird studies in the 90s. Taxonomic groups, however, ranged across the whole time period and did not cluster and as such bias the results (Fig. 6, colour code).

**Figure 6:**
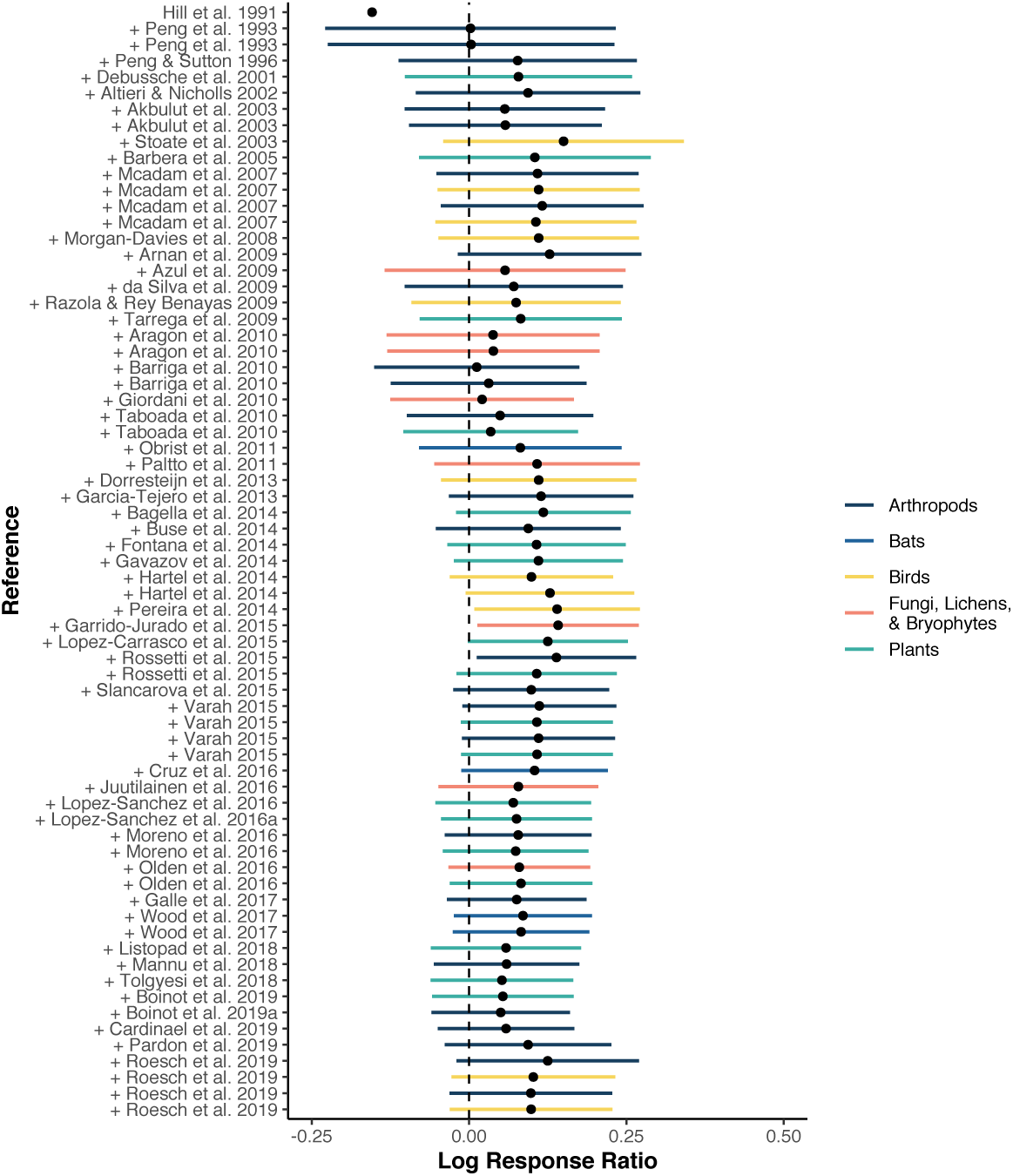
Cumulative forest plot, showing the summary effect sizes with always one individual effect size added over time. Colour code for the biodiversity groups indicate no clustering of any group in the time series.

## Discussion

European silvoarable systems host higher biodiversity than conventional agriculture, but show a tendency towards lower diversity than forests. In silvopastoral systems there was no evident benefit over either single-use system. Abandoning traditional agroforestry systems and leaving them to shrub encroachment and natural succession did not increase or reduce biodiversity systematically, such as suggested in other studies and is likely to depend on the number of years they were left abandoned [52, 53]. Birds and arthropods were the taxonomic groups with significantly higher diversity in an agroforestry system. The higher diversity of arthropods in agroforestry could not be traced back to any particular subgroup such as beetles, spiders or bees. Even within the taxonomic subgroups effects were heterogeneous. Spider diversity, for example, was found to be higher in agroforestry compared to a forest in one study [54, moderate-evidence study], but showed the opposite effect in another study [55, weak-evidence study].

Agroforestry covers around 10% of the agricultural area in the European Union [15]. Among them are traditional and very long established agroforestry sites, such as the Mediterranean Dehesas and Montado, traditional Spanish and Portugese silvopastures [3]. Land-use history, i.e. the age of the agroforestry system and the previous land-use type, may have a strong impact which is hardly reported or even known by the primary-study authors [20]. As such an older agroforestry system may harbour a different biodiversity than a newly established one; and the same holds for an old-grown forest relative to a more intensively managed younger forest site.

Additional unmeasured drivers operating at the landscape scale may equally determine the biodiversity. The implementation of agroforestry at the field scale does not guarantee the viability of populations of tree-dependent species, but could host these species if additional forest patches are found nearby [56, 57]. Invertebrates for example profit from a diverse landscape beyond the field scale [58, 59]. Our conclusion are largely based on species richness comparison; communities may well differ in their composition beyond richness [compare e.g. 22, 60, 61].

### Robustness of meta-analytical results

Meta-analysis of systematically searched literature provides evidence that is stronger than individual studies, unsystematic literature searches and qualitative synthesis [28, 62, 63]. Conclusion drawn from a meta-analysis nevertheless depend on the robustness of the result, i.e. whether minor changes, such as alternating the weighting, could reverse the conclusion. Weighting of studies traditionally occurs by inverse variance without considering the differences in study quality and design. In previous work, the underlying evidence and thus the reliability of individual study results has shown to be distinct depending on their study design [64, 65]. Weighting studies proportional to the evidence underlying each individual study is an alternative to the traditional weighting. In our case, results did not change with the alternative weighting approach, but can confirm the robustness of our conclusions.

Meta-analysis has established in ecology and as such updates of already existing meta-analyses can show how and whether conclusions may change over time. In a cumulative meta-analysis, adding new studies according to their publication date, we did not observe a declining effect as observed in other meta-analyses, but the effect remained stable despite very heterogeneous individual study results [66]. In our study we also found that other environmental variables have an influence on the agroforestry-biodiversity relationship. Meta-analysis builds on what is found in the literature, and additional categorical environmental variables used as moderators in meta-analytical models are rarely balanced. The results of our analysis is robust over time and adding new studies is unlikely to impact the results [67], but systematically adding studies on silvoarable systems, which in the current meta-analysis make up one third of silvopastoral-study contribution, could well influence the results. An increasing number of silvoarable studies may drag the overall effect size further towards the positive end and eventually turn the combined result to be significantly positive. Given that silvopastoral systems are dominant in Europe, we are nevertheless convinced that the ratio of silvopastoral and silvoarable studies in our meta-analysis reflects the proportion in which agroforestry systems in Europe occur and provide representative results [15].

Reproducibility of results is a sign of robustness, but challenging and often frail [68, 69]. The present meta-analysis and the analysis from Torralba et al. [23] have resulted in different conclusions as we failed to reproduce their results. While Torralba et al. [23] concluded that agroforestry has a positive effect on biodiversity in general, we could confirm a benefit only in relation to conventional agriculture. A possible explanation is the different set of studies used in their meta-analysis. Their definition of agroforestry includes studies on hedgerows and woody riparian buffers bordering agricultural field, which we did not consider as agroforestry as they are not actually under silvicultural use. They have also missed study results from biodiversity studies that reported disadvantages of agroforestry [e.g. 70, 71]. Successfully consolidating different results could be achieved by clearly communicating the context in which they apply, providing code and data used in the analysis to posthoc identify differences, and a ranking scale communicating, how confident scientists are with their statements. This is desirable to support decision makers, and has been demonstrated for the policy-relevant IPCC reports [28, 72, 73]. In this specific case, where reviews with the same attempt on similar data yield different results, such a confidence statement may indicate that both reviews are indeed very similar in their assessment. In a subgroup analysis of Torralba et al. [23] distinguishing between fungi, arthropods, plants and birds, only birds were significantly positive, which we could confirm in our analysis. In contrast to their results, we have to emphasize that results are heterogeneous. Our review suggests weak effects, and we are only moderately confident about these findings, supposing that the main driver for biodiversity cannot be found in agroforestry but may lie at the landscape scale or be dependent on land-use history.

## Conclusion

Agroforestry increases biodiversity in silvoarable systems compared to agriculture and in general for birds and arthropods, but benefits were small and there was no overall positive effect of agroforestry on biodiversity. Outcomes were influenced by the heterogeneity of effect sizes and silvopastoral systems did not show a benefit over either single-use system. While previous reviews were enthusiastic and considered agroforestry to have led to an increase in biodiversity [23], we need to call for caution. In the present evidence assessment, we have identified only few studies providing results based on strong evidence and have found a heterogeneous picture, suggesting other variables to interact with positive or negative effects from agroforestry. Systematic reviews and meta-analyses are providing the best available evidence, but they do not automatically guarantee reproducibility. They depend on the quality, quantity and comparability of studies used in the analysis. We suggest to resolve these issues by a detailed reporting (1), data provision (2) and the communication of heterogeneity (3). Our study provides results embedded in the context in which agroforestry can lead to a benefit for biodiversity. The use of these results can enrich the discussion on how future subsidies from the Common Agricultural Policy of the European Union can further incorporate agroforestry measures. Future studies on landscape parameters and land-use history are required to disentangle the context in which agroforestry is beneficial for biodiversity.

## Ethics approval and consent to participate

Not applicable.

## Consent for publication

Not applicable.

## Availability of data and materials

All data required for the replication of the analysis are available in the supplementary material.

## Competing interests

The authors declare that they have no competing interests.

## Funding

This work was initially supported by the 7^th^ framework programme of the European Commission in the project ‘Operational Potential of Ecosystem Research Applications’ (OPERAs, grant number 308393, www.operas-project.eu). The first author was funded by the STAY! scholarship of the Neue Universitätsstiftung Freiburg. The article processing charge was funded by the Baden-Württemberg Ministry of Science, Research and Art and the University of Freiburg in the funding programme Open Access Publishing.

## Authors’ contributions

ACM and CFD designed the study. ACM and MK collected the data. ACM analysed the evidence base. ACM performed the meta-analysis. ACM and CFD wrote the manuscript.

## Acknowledgements

We thank authors of primary studies used in this article to contribute to our meta-analysis by sharing their data and providing additional information to their publications. We also want to thank Amelie Göbel who has initiated this project with her master thesis and provided the evidence assessment of some of the primary studies. We are grateful to the initiative ‘Publication Partners during the Covid-19 lockdown’ and our publication partners Dominic Andreas Martin for the wonderful figure design and comments on earlier versions and Jonathan Spencer for text editing.

## Additional Files

Additional Files will be provided upon request.

## Notes

### Competing Interest Statement

The authors have declared no competing interest.

